# Working memory readout varies with frontal theta rhythms

**DOI:** 10.1101/2025.03.27.645781

**Authors:** Hio-Been Han, Scott L. Brincat, Timothy J. Buschman, Earl K. Miller

## Abstract

Increasing evidence suggests that attention varies rhythmically, phase-locked to ongoing cortical oscillations. Here, we report that the phase of theta oscillations (3–6 Hz) in the frontal eye field (FEF) is associated with temporal and spatial variation of the read-out of information from working memory (WM). Non-human primates were briefly shown a sample array of colored squares. A short time later, they viewed a test array and were rewarded for identifying which square changed color (the target). Better performance (accuracy and reaction time) varied systematically with the phase of local field potential (LFP) theta at the time of test array onset as well as the target’s location. This is consistent with theta “scanning” across the FEF and thus visual space from top to bottom. Theta was coupled, on opposing phases, to both spiking and beta (12–20 Hz). These results could be explained by a wave of activity that moves across the FEF, modulating the readout of information from WM.

## Introduction

Cortical activity fluctuates rhythmically, which has consequences for its function. This begins with sensory systems that sample the external world with periodicity^1,2^. It is also evident in visual attention. Even when trained to sustain steady visual attention on a single location, attention nonetheless fluctuates. The ability of non-human primates (NHPs) to detect a target at that location waxes and wanes on 3–6 Hz (theta) cycle^3^. These alternations of better and worse performance align with the phase of theta local field potential (LFP) oscillations in the frontal cortex. This periodicity of perceptual and attentional processes raises the possibility that other cortical functions, not just those involved in selecting and processing external inputs, might synchronize to particular cortical rhythms.

We examined working memory (WM), which is linked with attention^4–6^. Neural spiking during WM retention shows burstiness that co-varies with local field potential (LFP) rhythms across a wide range of frequencies^7–13^. But it is not known if WM function per se cycles at a base frequency, like attention does in theta. Thus, we sought to test whether there was a rhythmic component to WM-dependent behavior.

We analyzed neural activity recorded from the frontal eye field (FEF) in two non-human primates (NHPs) performing a change identification WM task (**Fig 1a**)^14,15^. NHPs were shown a sample array of colored squares (set size: 2–5), followed by an 800–1000 ms memory delay. Then a test array appeared in which one of the squares had changed color (the target). The NHPs made a direct saccade toward the target to receive a juice reward.

**Figure 1.**
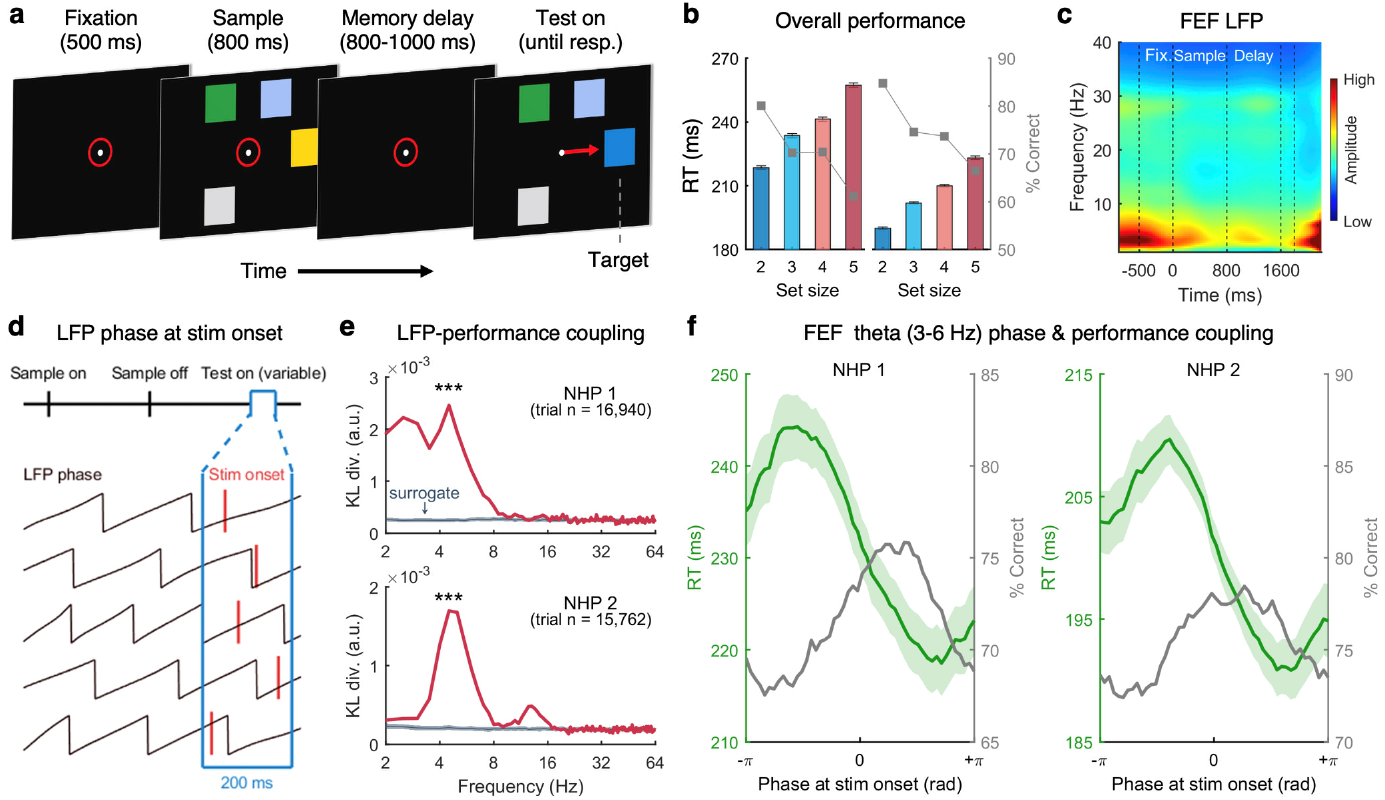
FEF theta modulates WM task performance. (**a**) WM task sequence. NHPs indicated which square changed color (the target) by making a saccade to its location. Timing of the test array was randomly determined because of the variable length of memory delay period. (**b**) Performance of two NHPs as a function of the number of items held in WM (colored bar graphs: RT, Gray squares: % Correct). (**c**) Oscillatory amplitude (grand-averaged) across time course of WM task. (**d**) Schematic illustration of the LFP phase at the timing of test stimulus onset. Across all trials, LFP phase was roughly uniformly distributed at test array onset (**Fig S2**). (**f**) Non-uniformity of RT distribution over LFP phase measured by KL-divergence in FEF (top: NHP 1, bottom: NHP 2). Non-uniformity of the distribution of RTs peaked at 4–5 Hz in FEF. Surrogate data were obtained by trial-shuffled bootstrapping (1,000 samplings). (**e**) Behavioral performance (green: RT, gray: % Correct) as a function of FEF theta phase for NHP 1 (left) and NHP 2 (right). Shaded area shows 1 SEM. *** *p* < .001 for the result of Wilcoxon’s signed-rank test.

WM performance depended on both the FEF theta phase at test array onset and the target’s position. The findings suggest a traveling wave of activity across the FEF, leading to a top-to-bottom spatial sampling that influenced WM readout.

## Results

### Behavioral performance cycled with FEF theta phase

Each NHP completed 14 recording sessions (NHP 1: 16,940 trials in total; NHP 2: 15,762 trials in total). Behavioral performance declined as a function of set size, consistent with the limited capacity of WM (**Fig 1b**)^14,16^. LFPs and spiking activity were recorded in FEF during task performance. Overall LFP power was characterized by prominent theta oscillations extending through the trial (**Fig 1c**).

To determine if behavior varied with LFP, we determined the instantaneous phase for frequencies from 2–64 Hz at the time of the test array onset. Note that due to the unpredictable time of the test array (**Fig 1d**), LFP phase in FEF at test array onset was not time-locked to external events. LFP phase was compared to the NHP’s reaction time (RT) and accuracy (percent correct change identification) by measuring the Kullback-Leibler (KL) distance from a circular uniform distribution across LFP phases, separately at each frequency. Large KL distances indicate performance reliably differs between LFP phases at test array onset.

This revealed a relationship between FEF theta and task performance. There was a clear peak within the theta range (∼5 Hz) for both NHPs, indicating a correspondence between behavior and FEF theta phase when the test array appeared (**Fig 1e**). Behavioral performance—both RT and accuracy—showed significant modulation by the phase of FEF theta (3–6 Hz; *p*s < .001 for RT and accuracy, for both NHPs). RT was faster and accuracy was higher if the test array appeared during the falling (“good”) phases (i.e., 0 to +π rad) of theta relative to the rising (“poor”) phases -π to 0 rad) of theta (**Fig 1f**). This theta modulation was stronger in trials with higher WM load (**Supplemental Fig S1**). We confirmed that LFP phase at test array onset was not phase-locked to trial events and thus uniformly distributed across trials (see **Supplemental Fig S2** for event-related potentials and inter-trial phase coherence). This confirmed that behavior was influenced by the FEF theta phase at test array onset.

### Theta sampled visual space sequentially

We found that behavior not only depended on the theta phase at test array onset, it also depended on the location of the target. Errors were not randomly distributed in space. Rather, incorrect choices tended to be near the correct target. **Figure 2b** shows a distribution of choices when the target was at the 3 o’clock position. When they chose the wrong array item, that item was more likely to be at 1 o’clock and 5 o’clock compared to further locations (e.g., 9 o’clock). **Figure 2c** shows the choice distribution after rotating and aligning saccadic landing positions relative to the target (for results of each location, see **Supplemental Fig S3**). The proximity of items to the target location and the frequency of choice errors showed a negative correlation (**Fig 2d**, NHP 1, *r* = –0.70, *p* < .01; NHP 2, *r* = –0.65, *p* < .01). We found a relationship between this spatial choice bias and theta phase .

**Figure 2.**
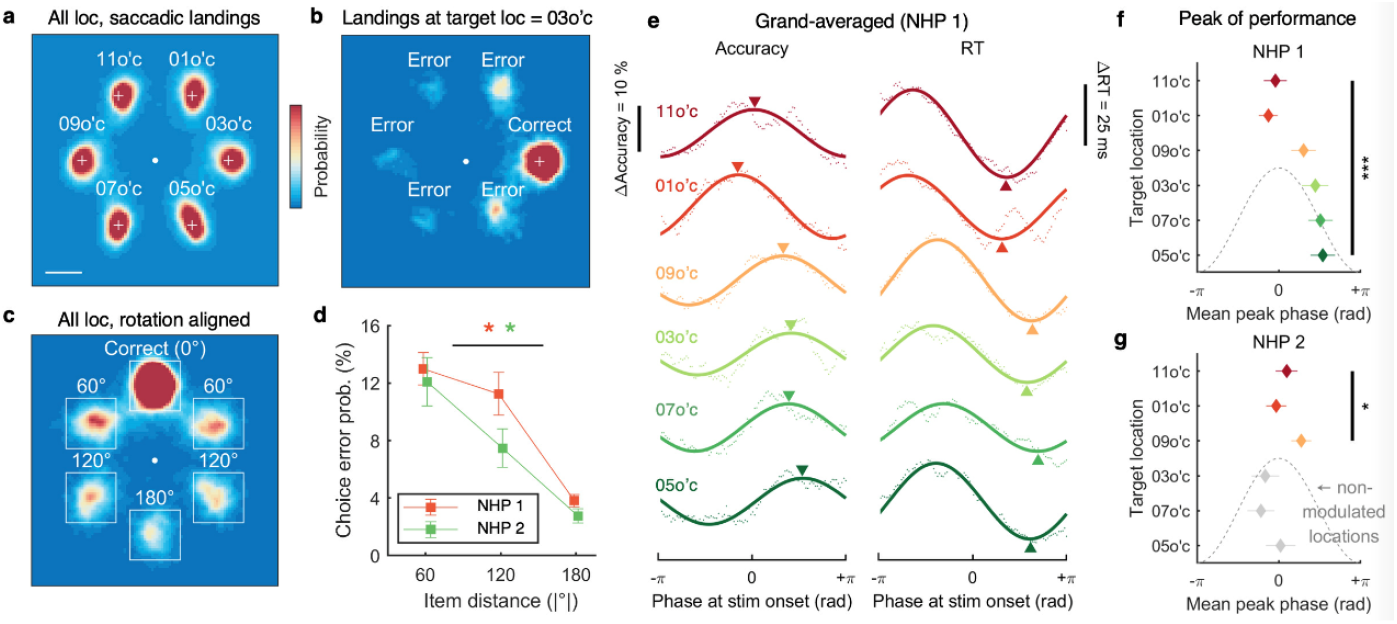
Retinotopic location-dependent sequential sampling of WM items. (**a**) Two-dimensional histogram of saccadic landing position for all possible target locations (all trials from two NHPs aggregated, trial n = 32,702). (**b**) Saccadic landing histogram for the trials with target location at 3 o’c.**(c)** Saccadic landing histogram for all possible target locations, rotated to align target location at 0° (top).**(d)** Monotonic decrease of choice errors as a function of item distance from target. * *p* < .01 for Pearson’s correlation coefficient. (**e**) Fluctuation of WM task retrieval performance (left: percent correct, right: RT) as a function of FEF theta phase at test display onset, for 6 possible target locations for NHP 1. Solid lines show sinusoidal curve fits to accuracy (dots) across theta phase. Colored triangles denote the optimal phase from the sinusoidal fits. (**f**) Peak phase of ‘good’ theta phase at each target location estimated from single session-level curve fitting. (**g**) Same as (**f**), but for NHP 2. Gray-colored locations did not show theta-rhythmic modulation. * *p* < .05, ** *p* < .001 for the result of circular ANOVA test in (**f**-**g**). Error bars indicate 1 SEM.

The good and poor phases of theta seemed to sample visual space sequentially from the top to bottom of the array. This was revealed by an analysis of performance as a function of theta phase and position of the target in the array. We fitted a sinusoidal function to the distribution of performance (accuracy, RT) over FEF theta phase at test array onset, separately for each target location. For both NHPs, a circular ANOVA showed significant differences in the peak performance phase across locations (*p* < .001 for NHP 1, *p* < .05 for NHP 2. For NHP 1, theta phase modulated performance at all six array locations in a systematic fashion. Performance was better for the top two array locations (11 o’clock and 1 o’clock) when the array appeared at or near the peak of FEF theta. At lower locations (9 o’clock and 3 o’clock, then 7 o’clock and 5 o’clock), performance peaked when the test array appeared at progressively later phases of theta (**Fig 2ef, NHP 1**). NHP 2 showed significant theta modulation at three out of six locations (**Fig 2g** and **Supplemental Fig S4a**). This difference between NHPs may stem from their different task strategies. NHP 1 performed equally well across all target locations (*F*(5,78) = 2.210, *p* = .062). In contrast, NHP 2 showed uneven performance across locations with higher accuracy at the lower location that did not show sequential sampling (*F*(5,78) = 14.312, *p* < .000, **Supplemental Fig S4**). Thus, NHP 1 seemed to split processing evenly across all locations (and thus sampled all locations). NHP 2 instead focused mainly on the lower locations and thus did not “scan” the full visual space like NHP 1. Thus, we observed sequential theta modulation at visual field locations where NHPs divided their attention.

### Neural information cycled with theta

Theta was composed of alternating excitatory and inhibitory cortical states (**Fig 3**). Phase-amplitude coupling revealed that theta phase modulated beta (12–20 Hz) power. Beta power was lowest during the rising theta phases and higher during the falling theta phases (**Fig 3a**). This was significant for both the sample and memory delay (**Fig 3b**, *p*s < .01). Spiking was also coupled to theta (**Fig 3c**, *p*s < .01). Spike rate was highest at the troughs of theta (near -π and +π, (Fig 3d). This was when beta was lowest (around -π) and late in the falling theta phase (around) +π). Spike rate was lowest just after the peak of theta when beta power was highest (**Fig 3d**).

**Figure 3.**
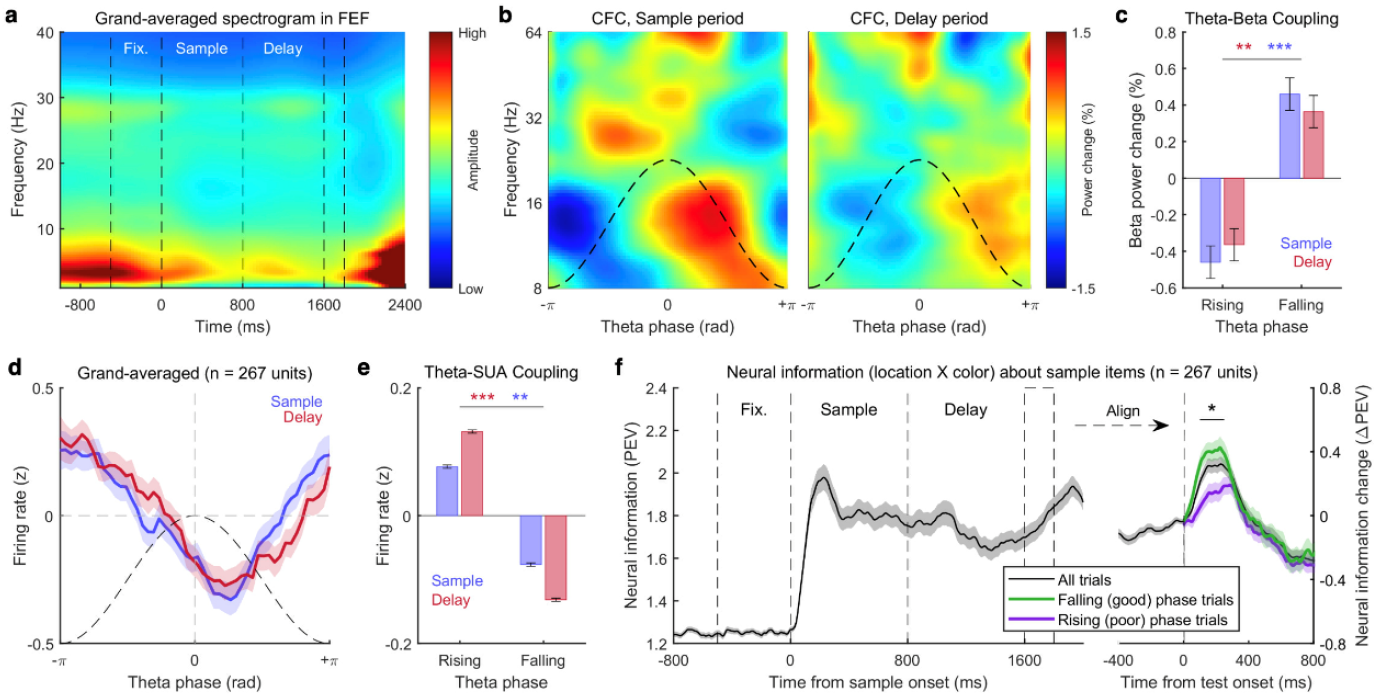
Theta rhythmically modulates beta power and SUA in FEF. (**a**) Cross-frequency coupling between theta phase and high-frequency oscillation power during sample (0–800 ms, left) and delay period (800–1600 ms, right). Dashed lines denote one cycle of theta. (**b**) Beta power comparison between rising (behaviorally “poor”) and falling (“good”) phase of theta. (**c**) Firing rate comparison between rising and falling phase of theta. (**d**) Grand-averaged SUA as a function of theta phase (delay period only). (**e**) Time course of neural information contained in SUA calculated by PEV, time-aligned by sample array onset (left) and test array onset (right). Shaded area and error bars indicate 1 SEM. * *p* < .05, ** *p* < .01, *** *p* < .001 for the result of Wilcoxon’s signed-rank test.

We measured neural information about WM items (i.e., the sample array) using FEF spiking activity to examine its interaction with theta phase. Using a GLM, we predicted item location and color, quantifying explanatory power with percent explained variance (PEV). Figure 3e shows that neural information increased after sample onset and remained stable during the delay period, as expected. After test array onset, it increased again, suggesting WM retrieval triggered by the test array. The effect of theta phase in the memory delay emerged at test array onset. More information was present when the test array appeared during the falling (good) phase of theta than during the rising (poor) phase (**Fig 3e**). The good vs. poor phases were defined by the instantaneous theta phase at test array onset (**Fig 1f**). Visual information arrived in the FEF with a latency of about 100 ms (see ‘sample’ period in **Fig 3e**). Thus, information about the test array would arrive, not during the instantaneous (inhibitory) phase but in the opposite excitatory phase.

## Discussion

Our results indicate that the phase of frontal theta during a memory delay was associated with the readout of information from WM. The ability to detect a change (a target) in a visual scene (an array of items) from a similar scene held in WM fluctuated with theta oscillations in the FEF. Performance depended on both the theta phase when the comparison scene appeared as well as the location of the target. It appeared as if a theta wave was “scanning” the WM representation of the scene from top to bottom. This could be explained by a traveling wave moving across a retinotopic FEF. Performance improved when the theta wave happened to align with the location of the target when the comparison scene appeared. Cortical excitatory/inhibitory states and neural information also cycled with frontal theta, suggesting a possible mechanism for its effects on behavior.

Previous work has shown that selective attention waxes and wanes in theta in correspondence with frontal theta rhythms. During periods when attention is ostensibly sustained at a constant location in space, behavioral performance varies with a temporal periodicity of ∼4–5 Hz^17,18^. Studies of human EEG/ECoG^19–21^ and NHP LFPs^22–24^ have shown this rhythmicity reflects ongoing theta oscillations in frontoparietal cortex. Theta oscillations are often induced when attention is directed toward one of multiple competing stimuli^25,26^. Depending on the phase of these theta oscillations when a probe stimulus is shown, behavioral performance can vary dramatically^3,19–22,24^. These results show that ongoing cortical theta oscillations can modulate attention to external sensory inputs.

Our results indicate that *internal* WM representations also cycle with frontal theta. This is consistent with human behavior^27–29^ and EEG^30,31^ studies. In our case, the theta cycling modulated the behavioral readout of WM. Our results share many commonalities with the attention literature. Prior work on attention also found the strongest modulation within a similar 4–5 Hz band^20–22,24^. Attention studies have likewise found behavior is optimal when probed during the falling phase of frontal theta^22^. Prior work also showed similar theta modulation of spiking and higher-frequency activity^21,22,24^. Overall, similarities between our results and previous studies of attention strongly suggest a common mechanism may be at play. We propose it reflects shared control mechanisms deployed for both attention and WM. This is consistent with many previous proposals suggesting shared control of attention and WM^3–6^.

Our results also build on previous work by demonstrating that theta modulation has an orderly structure across visual space. Previous studies have typically contrasted single locations inside and outside the focus of attention. Their results have been interpreted as “good” and “poor” theta phases alternating at the attended location, while “poor” and “good” theta phases oscillate in anti-phase at the unattended location^3,32^. This can be equivalently thought of as “good” and “poor” theta phases alternating back and forth between the attended and unattended locations. Our study generalizes this idea to a structured shift of theta phases when resources are not focused on a single location but divided across visual space. The optimal theta phase for behavior varied by retinotopic target location, progressing from the top to the bottom of the visual field.

This could be explained by a traveling wave of activity across the cortical surface during the memory delay. Traveling waves have been observed in a number of cortical areas, suggesting they may be a ubiquitous motif of cortical processing^33–38^. Our results would suggest a wave sweeping across the polar angle dimension of the FEF topographic map, arrayed along the anterior-posterior axis^39^. In fact, waves of theta oscillations propagating in the posterior-to-anterior direction have been observed in human frontal cortex^40^.

A simple explanation of the results is that a traveling wave of excitation enhances processing when it aligns with the target. The behaviorally good phase was the inhibitory phase (when spiking and gamma is falling and alpha/beta is higher). However, this was the *instantaneous* phase when the test array appeared. Spiking activity indicates that information reaches the FEF around 100–200 ms after onset (see ‘sample’ period in **Fig 3e**), matching the latency of effects following test array onset (**Fig 3e**). This suggests that test array information in spiking arrives in the FEF not during the instantaneous inhibitory phase, but during the following excitatory phase of theta (i.e., when spiking and gamma is rising and alpha/beta is lower). Thus, the excitatory phase was the good phase in terms of brain mechanisms. On the other hand, it is unclear exactly when this input becomes functionally relevant—it may instead arrive during an inhibitory phase. If so, theta could play a role in top-down WM processes such as stabilizing internal representations to reduce interference through beta. Beta, which was higher during the inhibitory theta phase, has been associated with stabilizing cortical representations^41^ and top-down control of sensory processing^19,21,42^. In either case, our results suggest theta wave modulates how information is read out from WM.

Many theories have emphasized the role of theta as a temporal framework for structuring cognitive processes. The *rhythmic theory of attention*^3^ proposes visual attention sequentially samples perceptual inputs within a theta cycle. The *theta-gamma neural code*^10^ suggests that within a single theta cycle, distinct neural ensembles encoding different information are activated in succession across several nested gamma cycles, enabling multiplexed representation. More recently, the *rhythmic attentional scanning* model^43^ suggested each theta cycle acts as a selection window, determining which of multiple competing representations is propagated downstream. These frameworks collectively suggest that theta actively segments cognitive processing into periodic sampling windows, which our results support.

Our results also suggest that theta modulation plays a key role under high cognitive demands, such as when memory load increases and resources are divided across multiple items or locations—a characteristic trait of theta oscillations^25,26,44–47^. Frontal theta, involved in active resource control^48^, is known to increase with cognitive load—much like a car engine straining uphill^49^. In visual attention studies, theta has been linked to the intermittent sampling of unattended locations^24^. This may explain why theta modulation is often stronger outside the primary focus of attention^24^. Sustained attention to a single location likely involves continuous resource allocation, reducing the need for theta-driven sampling. In our task, there was no primary focus. WM resources and readout were meant to be evenly distributed. The NHPs varied in how well they achieved this. We observed stronger theta modulation at scene locations where behavior indicated resource division. This theta mechanism may generalize to any context where the brain must manage multiple simultaneous representations, whether external or internal.

Our findings demonstrate that WM readout varies with frontal theta oscillations, with behavioral performance depending on FEF theta phase. Theta appeared to structure the spatial organization of WM, with retrieval performance varying systematically across retinotopic space. Our findings provide further evidence that cognition is intrinsically linked to cortical oscillatory dynamics.

## Methods

All procedures followed the guidelines of the Massachusetts Institute of Technology Committee on Animal Care and the National Institutes of Health.

### Experimental Model and Subject Details

One adult rhesus macaque (*Macaca mulatta*, NHP 1: male, 13 kg) and one adult male cynomolgus monkey (NHP 2: male, 6 kg) were trained to perform the task. For neural recording, multiple epoxy-insulated tungsten electrodes (FHC, Bowdoin, ME USA 04287) were inserted using custom-built manual screw microdrives. FEF was targeted via co-registration of structural MRI scans with standard atlases, and confirmed by microstimulation-driven saccades. The electrodes were acutely lowered at the beginning of every recording session (*n* = 14 sessions for each NHP) and settled for at least 2 hours before recording, then retracted after it. For further details on surgical procedures and animal handling, please see our previous publication with the same dataset.

### Methods details

#### Behavioral protocol and data acquisition

The behavioral paradigm was controlled with the MonkeyLogic program^50–52^. Each trial began with a 500 ms fixation period, followed by an 800 ms sample period where an array of colored squares was displayed. After a variable memory delay period (800–1000 ms), a test array appeared, identical to the sample array except for a color change in one randomly selected target. NHPs were required to make a single saccade to the changed item. Eye movements were tracked using an ISCAN infrared system (240 Hz) throughout all sessions. Stimuli were 1° colored squares, with two possible colors (color A, and color B) at each location, randomized daily to prevent long-term memorization. Six item locations (roughly 1, 3, 5, 7, 9, and 11 o’clock) were used each session that were ±75 angular degrees from the horizontal meridian and 4° to 6° from the fixation point. For the analysis, gaze coordinates at saccading landings on six target locations were translated to ensure symmetry and then rotated so that the target location was positioned at 0 degrees (top of the screen). NHPs completed at least 720 correct trials per session. Invalid trials (e.g., failure to fixate before test array onset) were excluded (trial survival rate: 76.29% for NHP 1, 71.09% for NHP 2). As a result, total 16,940 trials for NHP 1 (load 2: 29.56%, load 3: 28.56%, load 4: 21.46%, load 5: 20.43%) and total 15,762 trials for NHP 2 (load 2: 27.44%, load 3: 26.95%, load 4: 20.21%, load 5: 18.44%) were included. For more details on the behavioral paradigm, please see our previous publication that used the same dataset^14^.

#### LFP data acquisition and preprocessing

Continuous LFP data was amplified, band-passed filtered (3.3–88 Hz), and digitized at 1 kHz sampling rate (Plexon Multi-channels Acquisition Processor). All signals were referenced to ground. Any 60 Hz line noise, 85 Hz noise related to the monitor refresh rate, and their harmonics were estimated and removed offline using an adaptive sinusoid fit method. For all LFP analyses, LFPs were averaged across all simultaneously recorded FEF electrodes (*n* = 7.93 ± 2.00 per session) to estimate a representative LFP signal for FEF.

To estimate theta phase at the timing of test array onset, LFP was band-pass filtered at 3–6 Hz with zero-phase FIR filters (MATLAB’s *filtfilt*.*m* function). After band-pass filtering, the Hilbert transform was used to obtain instantaneous angle (MATLAB’s *hilbert*.*m* and *angle*.*m* functions). Theta phase was divided into discrete phase bins (50 bins) for further analyses. The strength of coupling between LFP phase and behavioral performance was measured by the Kullback-Leibler divergence between the observed histogram of the behavioral measure (RT or accuracy) and a circular uniform distribution.

To obtain amplitude (power) spectrograms, a fast Fourier transform was applied with a sliding Hanning window (window size = 1024, 100 ms step). For cross-frequency coupling analysis, the LFP signal was narrow-band-filtered (0.5 Hz step with 2 Hz bandwidth for 8–16 Hz; 1 Hz step with 4 Hz bandwidth for 16–32 Hz; 2 Hz step with 8 Hz bandwidth for 32–64 Hz) using Butterworth 5^th^ order filter.

#### Spike data analysis

Spikes were sorted into isolated single units manually using waveform features (Plexon Offline Sorter), as previously described^14^. Then, two criteria were applied for excluding units from the analysis: (1) units with less than 30 trials sampled for each condition were excluded; (2) units with a firing rate less than 1 Hz during the task period (–0.5 to +1.6 s relative to sample stimulus onset) were excluded. As a result of applying these two criteria, 76.19% of units (leaving *n* = 267 units) from FEF were included in the analysis.

To investigate how spiking data encodes WM information (location x color), we analyzed individual SUA firing rates using a general linear model (GLM) and calculated its effect size (i.e., PEV measured by η ^2^, which quantifies how much of a unit’s firing rate variability is attributed to stimulus location and identity). Since the color of each object was unknown to the NHP before sample array onset, we assessed how well SUA represented this information at all time points. PEV was computed using a single ANOVA model, where all locations and item identities were included in a unified model with dummy-coded variables representing location × item identity: y = Σ bX where X = 0, 1, 2, represented absent, color A, and color B, respectively, for each location. PEV was calculated using the η ^2^ formula and summed across all model variables: PEV = (1 / SS_Total) * Σ SS_X * 100. This approach provides a comprehensive assessment of how spiking activity encodes WM information across all spatial locations, resulting in a measure of the overall strength of WM representations.

### Quantification and Statistical Analysis

Pearson’s *r* was used for correlation analysis. For testing sample means (or medians) against a null value (appropriate for a one-sample *t*-test), non-parametric Wilcoxon’s signed rank test was used. Circular ANOVA was used when the dependent variable was angular phase data, implemented by CircStat toolbox^53^ in MATLAB (function *circ_wwtest*.*m*).

## Acknowledgements

This work was supported by the Picower institute for learning and memory, the Office of Naval Research MURI N00014-23-1-2768, NEI 1R01EY033430-01A1, Office of Naval Research N00014-22-1-2453, the Freedom Together Foundation, and National Research Foundation of Korea Grant (NRF-2022R1A2C3003901; RS-2024-00460928), and by the Research Program funded by the SeoulTech (Seoul National University of Science and Technology).

## Author contributions

Author Contributions: H.-B.H. and E.K.M. designed the study. T.J.B. and S.L.B. conducted the research. H.-B.H., S.L.B., and T.J.B. analyzed the data. H.-B.H., S.L.B., and E.K.M. wrote the manuscript, and E.K.M. supervised the study.

## Resource availability

Requests for further information and resources should be directed to and will be fulfilled by the lead contact, Earl K. Miller.

