## Supplemental for "Working memory readout varies with frontal theta rhythms"

List of contents

Supplementary Figure S1
Supplementary Figure S2
Supplementary Figure S3
Supplementary Figure S4

### Supplementary Information

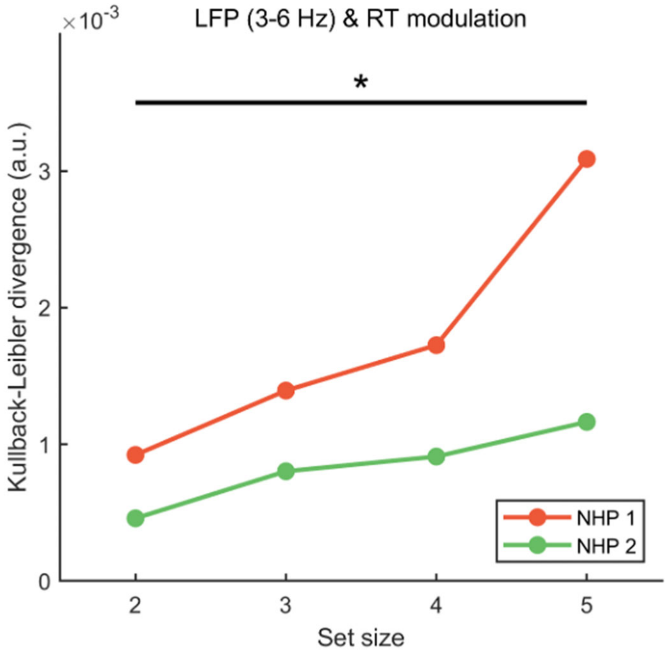


**Figure S1**. **Effect of task load on theta phase modulation of reaction time, related to Figure 1.** The higher WM load was associated with the stronger theta phase modulation (Pearson's $r$ = .663, $p$ = .036).

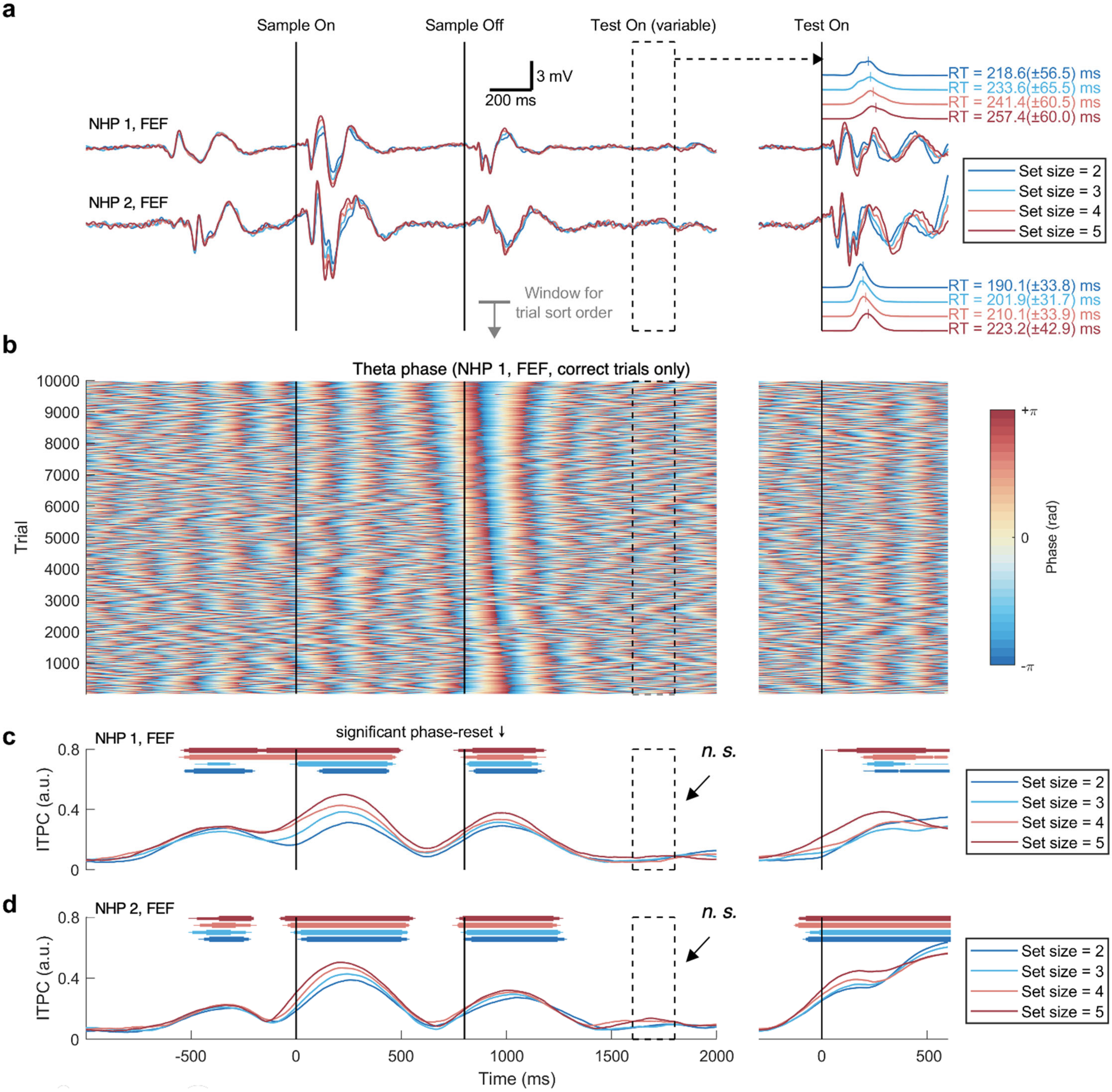


**Figure S2**. **Short-lasting phase-reset effect of visual event on FEF theta, related to Figure 1.** (**a**) Event-related potentials (ERPs) to visual stimuli in the frontal eye field (FEF) are shown, aligned for the sample array onset (left) and test array onset (right). There is no noticeable ERP component before the jittered test array onset timing (black dotted box). (**b**) Example single-trial phase traces, sorted by the mean phase value after the sample array offset (time window = 868-1018 ms, highlighted in gray solid line in (**a**)). (**c**) Inter-trial phase coherence (ITPC) calculated from NHP 1 FEF. Colored dots indicate statistical significance from t-test against the null hypothesis (i.e., no increase of ITPC from the pre-fixation period). Small dot: $p < .05$, medium dot: $p < .01$, large dot: $p < .001$. (**d**) Same as (**c**), but for NHP 2. Note that the phase-resetting effect of the visual event (i.e., sample off) lasted approximately 500 ms, resulting in a random phase distribution around the test array onset timing (black dotted box).

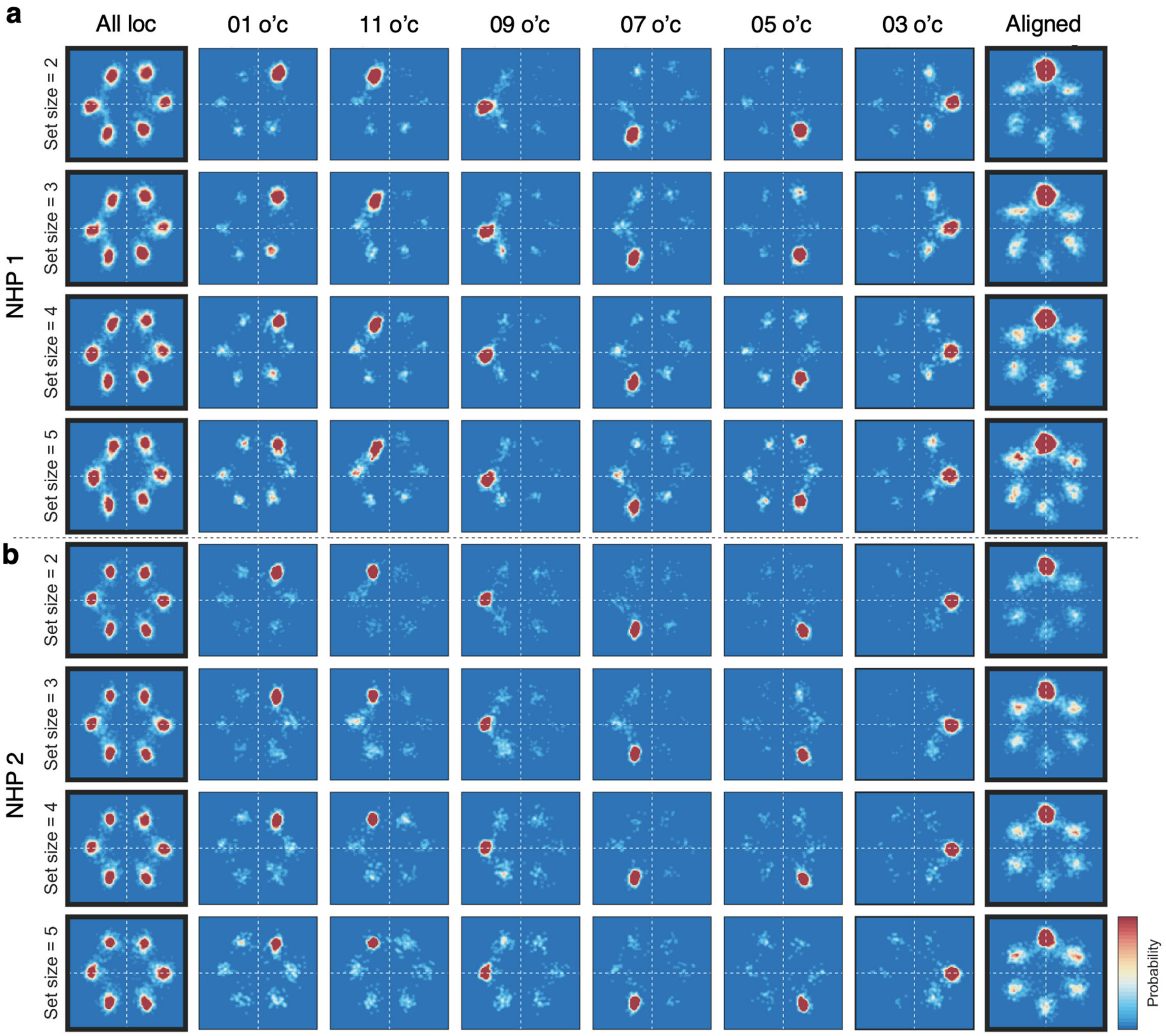


**Figure S3**. **Two-dimensional histogram of saccadic landing position for each target location, related to Figure 2.** (**a**) Saccadic landing position histograms from NHP 1 that show the spatial distribution of choice error, which shows false alarms were more likely localized near to the target location (rows: set size; column: target location). (**b**) Same as (**a**) but for NHP 2.

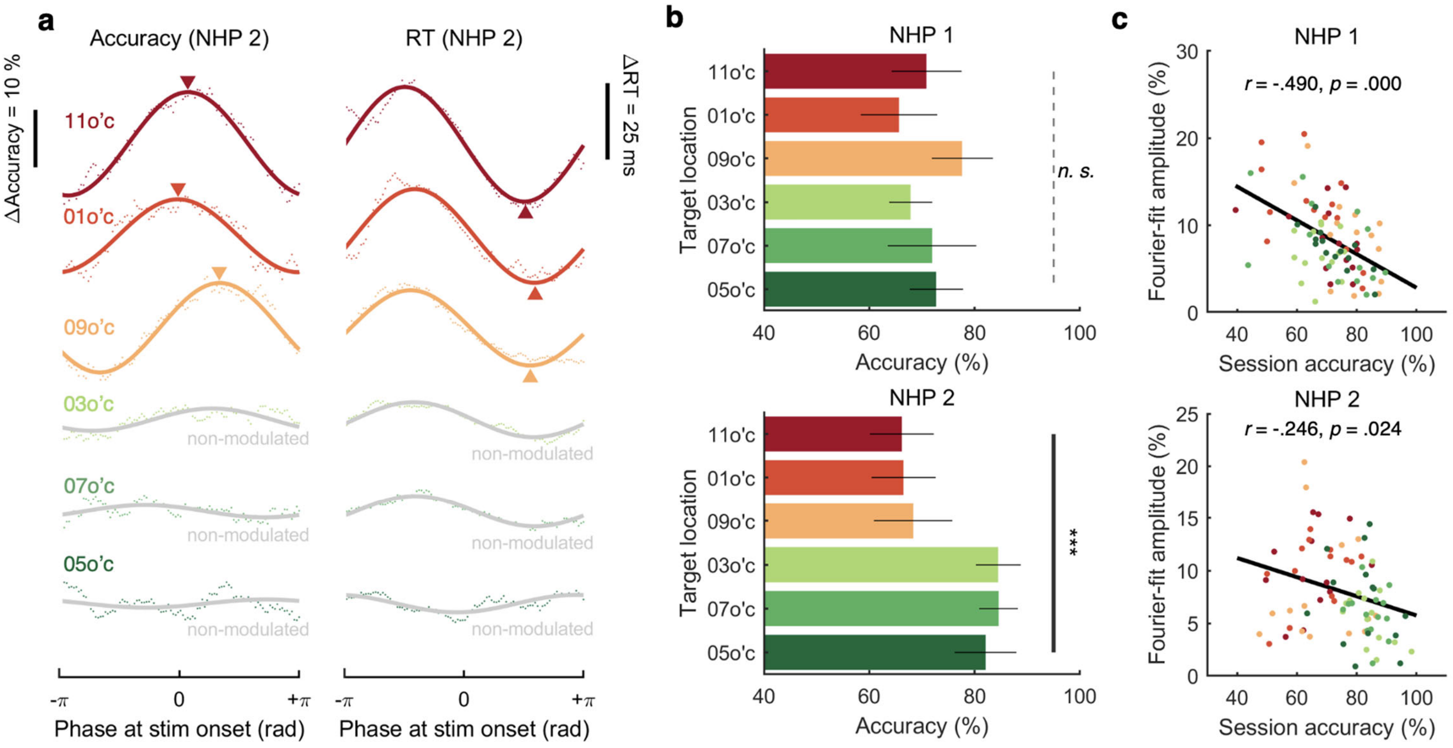


**Figure S4. Stronger FEF theta modulation is associated with lower task performance, related to Figure 3**. (**a**) Accuracy by target location for NHP 1 (top) and NHP 2 (bottom). For NHP 1, differences in accuracy across target locations were not statistically significant, ANOVA $F(5,78) = 2.210$, $p = 0.062$. In contrast, NHP 2 showed significant differences, ANOVA $F(5,78) = 14.312$, $p = 0.000$, suggesting the use of distinct task strategies for specific retinotopic locations (e.g., 3 o'c, 7 o'c, and 5 o'c). (**b**) Changes in task performance (accuracy) as a function of FEF theta phase at the onset of the test display for six possible target locations in NHP 2 (session N = 14, grand-averaged). Gray lines indicate target locations where sinusoidal curve fitting failed ('non-modulated' locations). (**c**) Negative correlation between daily accuracy and FEF theta-performance modulation. Each dot represents the accuracy for a given target location during a single session (N = 14 sessions per subject). The black solid line represents the least-squares polynomial fit. Pearson correlation tests were used for the statistical results in (**c**).
